# Discovery of Clioquinol and Analogues as Novel Inhibitors of Severe Acute Respiratory Syndrome Coronavirus 2 Infection, ACE2 and ACE2 - Spike Protein Interaction *In Vitro*

**DOI:** 10.1101/2020.08.14.250480

**Authors:** Omonike A. Olaleye, Manvir Kaur, Collins Onyenaka, Tolu Adebusuyi

## Abstract

Severe Acute Respiratory Syndrome Coronavirus 2 (SARS-CoV-2), the etiological agent for coronavirus disease 2019 (COVID-19), has emerged as an ongoing global pandemic. Presently, there are no clinically approved vaccines nor drugs for COVID-19. Hence, there is an urgent need to accelerate the development of effective antivirals. Here in, we discovered Clioquinol (5-chloro-7-iodo-8-quinolinol (CLQ)), a FDA approved drug and two of its analogues (7-bromo-5-chloro-8-hydroxyquinoline (CLBQ14); and 5, 7-Dichloro-8-hydroxyquinoline (CLCQ)) as potent inhibitors of SARS-CoV-2 infection induced cytopathic effect *in vitro*. In addition, all three compounds showed potent anti-exopeptidase activity against recombinant human angiotensin converting enzyme 2 (rhACE2) and inhibited the binding of rhACE2 with SARS-CoV-2 Spike (RBD) protein. CLQ displayed the highest potency in the low micromolar range, with its antiviral activity showing strong correlation with inhibition of rhACE2 and rhACE2-RBD interaction. Altogether, our findings provide a new mode of action and molecular target for CLQ and validates this pharmacophore as a promising lead series for clinical development of potential therapeutics for COVID-19.

## Introduction

Severe Acute Respiratory Syndrome Coronavirus 2 (SARS-CoV-2), a novel RNA betacoronavirus, is the causative agent for coronavirus disease 2019 (COVID-19), which has emerged as an ongoing global pandemic^1^. Worldwide, SARS-CoV-2 has spread rampantly to more than 188 countries/regions and has resulted in 18,847,261 confirmed cases, 11,390,018 recovered, including 708,540 deaths (https://coronavirus.jhu.edu/map.html). Within the United States alone, there are more than 4,825,742 cases, 1,577,851 recovered and a total of 158,300 deaths as of August 6^th^, 2020 according to the Johns Hopkins COVID-19 dashboard. About 80% of people infected with SARS-CoV-2 experience mild symptoms or are asymptomatic^2^; while a majority of symptomatic patients with moderate to severe symptoms have shown a broad range of clinical manifestation and/or significant complications, including severe pneumonia, multi-organ failure, acute cardiac injury, neurological damage, septic shock, acute respiratory distress syndrome (ARDS)^3–6^. Recent reports revealed that, individuals with pre-existing medical conditions have increased risk of COVID-19 related morbidity and mortality^7^. Currently, there are no U.S. Food and Drug Administration (FDA) approved drugs for the treatment of COVID-19; but several studies are investigating the potential utility of repurposing clinically approved drugs as treatment options for COVID-19^8–12^. To date, only Remdesivir, an inhibitor of RNA dependent RNA Polymerase has been granted emergency use authorization (EUA) for the treatment of hospitalized patients with severe cases of COVID-19^13^.

Historically, Clioquinol (5-chloro-7-iodo-8-quinolinol (CLQ)) and its derivatives belonging to the 8-hydroxyquinoline structural class, has shown potent broad-spectrum activity against clinically relevant pathogens^14–20^. More recently, CLQ and its analogues have been extensively investigated as potential treatments for cancer and neurodegenerative diseases^21–28^. Additional studies have also shown the involvement of CLQ in the efflux mechanisms of ATP binding cassette (ABC) transporters^29,30^ and the cellular autophagic pathway^31,32^, a critical process in the host defense machinery against viral infections^33^. Furthermore, using a high-throughput screen (HTS) and chemical genomics approach, Olaleye, O., et. al. identified and characterized CLQ and certain analogues as potent inhibitors of methionine aminopeptidase^17^, a universally conserved metalloprotease required for N-terminal methionine excision^34,35^. As an established metal chelator and zinc ionophore, CLQ modulates underlying molecular and physiologic machinery required for metal homeostatis^31, 32, 36–39^. Altogether, these pharmacologic properties of CLQ, makes it an attractive drug for potential targeting of Angiotensin Converting Enzyme 2 (ACE2).

ACE2 is a zinc metalloprotease and essential cellular receptor for SARS-CoV-2 entry into host cells^40–43^. Therefore, rapid identification of potent and selective ACE2 inhibitors have the prospects of accelerating the clinical development of preventative interventions and/or treatment options for COVID-19. ACE2 is mainly expressed in alveolar epithelial cells of the lungs, heart, kidney, and gastrointestinal tract^44,45^. Although, ACE2 is the cellular receptor for SARS-CoV-2 ^40–43^; ACE2 primarily functions as a carboxypeptidase that catalyzes the conversion of a single residue from angiotensin (Ang II), generating L-phenylalanine and Ang (1-7), a potent vasodilator, thus playing a critical role in controlling hypertension, renal disease, cardiac function and lung injury^46,47^. The crystalline structure of the ACE2 shows two domains; a N-terminal zinc metallopeptidase domain (MPD) capable of binding the viral envelope-anchored Spike (S) glycoprotein of coronaviruses, and a C terminal “collectrin-like” domain^48–50^. The interaction of the MPD of ACE2 and S glycoprotein of SARS-CoV-2 is the initial and critical step in viral infection by receptor recognition and fusion of host and viral cellular membranes^40–43^. In addition, viral entry requires priming of S protein by a host protease into S1 and S2 subunits, which are responsible for receptor attachment and membrane fusion, respectively^51–54^. A receptor-binding domain (RBD) of the S1 subunit specifically recognizes ACE2 on human cells^40–43^. Binding of the S1 subunit to ACE2 receptor triggers a conformational change in S glycoprotein from metastable pre-fusion state to stable post-fusion conformation, resulting in shedding of S1 and transition of the S2 subunit to expose a hydrophobic fusion peptide^42,55,56^. The initial priming at S1/S2 boundary promotes subsequent cleavage at the S2 site by host proteases, which is critical for membrane fusion and viral infectivity^54,57,58^. Therefore, targeting the interaction between human ACE2 receptor and the RBD in S protein of SARS-CoV-2 could serve as a promising approach for the development of effective entry inhibitors for potential prevention and/or treatment of COVID-19.

In this study, we evaluated the effect of CLQ, and two of its analogues (7-bromo-5-chloro-8-hydroxyquinoline (CLBQ14); and 5, 7-Dichloro-8-hydroxyquinoline (CLCQ)) on SARS-CoV-2 infection induced cytopathic effect (CPE) *in vitro*. In addition, we assessed the cytotoxicity of these compounds. Furthermore, we determined the impact of all three compounds on recombinant human ACE2 (rhACE2) interaction with the RBD on Spike protein of SARS-CoV-2; and independently assessed their effects on the exopeptidase activity of rhACE2. Here in, we discovered for the first time that CLQ, CLBQ14 and CLCQ effectively inhibits the novel SARS-CoV-2 infection induced CPE *in vitro*, inhibited rhACE2 and its interaction with Spike protein and rhACE2 exopeptidase activity in the low micromolar range. Thus, rapid optimization and pre-clinical development of CLQ and its congeners could potentially accelerate their consideration for re-purposing as potential antiviral agents against COVID19, first in non-human primate (NHP) models of SARS-CoV-2 infection, and subsequently in clinical trials.

## MATERIALS AND METHODS

### MATERIALS

#### Cell Growth Conditions and Medium

African Green Monkey Kidney Vero E6 cells (ATCC# CRL-1586, American Tissue Culture Type) were maintained using medium purchased from Gibco (modified eagle’s medium (MEM) Gibco (#11095); 10% fetal bovine serum (HI FBS) Gibco (#14000); Penicillin/Streptomycin (PS) Gibco (#15140); 10U/mL penicillin and 10μg/mL streptomycin (only in assay media)). For the SARS-CoV-2 infection induced cytopathic effect (CPE) assay, cells were grown in MEM/10% HI FBS and harvested in MEM/1% PS/supplemented with 2% HI FBS. Cells were batch inoculated with SARS-CoV-2 USA_WA1/2020 (M.O.I. ~ 0.002) which resulted in 5-10% cell viability 72 hours post infection.

#### Compounds and Preparation of Stock Solutions

The small molecule inhibitors, 5-chloro-7-iodo-8-quinolinol (Clioquinol, CLQ; C0187-Lot JJ01 SPGN), and 7-bromo-5-chloro-8-hydroxyquinoline (CLBQ14; B1190-P61JD-FD)); were purchased from TCI America. 5, 7-dichloro-8-hydroxyquinoline (CLCQ; D64600-Lot#STBH7389) and Zinc Chloride (ZnCl_2_; 208086-Lot#MKCL1763) were purchased from Sigma Aldrich. We prepared 10mM stocks solutions of the inhibitors in Dimethyl sulfoxide (DMSO; D8418-Lot#SHBL5613) purchased from Sigma Aldrich. For the CPE assay, compound samples were serially diluted 2-fold in DMSO nine times and screened in duplicates. Assay Ready Plates (ARPs; Corning 3764BC) pre-drugged with test compounds (90 nL sample in 100% DMSO per well dispensed using a Labcyte (ECHO 550) are prepared in the Biosafety Level-2 (BSL-2) laboratory by adding 5μL assay media to each well.

##### Method for measuring antiviral effect of CLQ, CLBQ14 and CLCQ

The SARS-CoV-2 infection induced cytopathic effect (CPE) assay and cytotoxicity assays were generated and performed through a sub-contract to Southern Research Institute (SRI), Birmingham, Alabama from Texas Southern University, Houston, Texas. The CPE reduction assay was conducted at SRI to screen for antiviral agents in high throughput screening (HTS) format as previously described^59,60^. Briefly, Vero E6 cells selected for expression of the SARS-CoV-2 receptor (ACE2; angiotensin-converting enzyme 2) are used for the CPE assay. Cells were grown in MEM/10% HI FBS supplemented and harvested in MEM/1% PS/ supplemented with 2% HI FBS. Cells were batch inoculated with SARS-CoV-2 (M.O.I. ~ 0.002) which resulted in 5% cell viability 72 hours post infection. Compound samples were serially diluted 2-fold in DMSO nine times and screened in duplicates. Assay Ready Plates (ARPs; Corning 3764 BC black-walled, clear bottom plates) pre-drugged with test compounds (90 nL sample in 100% DMSO per well dispensed using a Labcyte (ECHO 550) were prepared in the BSL-2 lab by adding 5μL assay media to each well. The plates were passed into the BSL-3 facility where a 25μL aliquot of virus innoculated cells (4000 Vero E6 cells/well) was added to each well in columns 3-22. The wells in columns 23-24 contained virus infected cells only (no compound treatment). Prior to virus infection, a 25μL aliquot of cells was added to columns 1-2 of each plate for the cell only (no virus) controls. After incubating plates at 37°C/5%CO_2_ and 90% humidity for 72 hours, 30μL of Cell Titer-Glo (Promega) was added to each well. Luminescence was read using a Perkin Elmer Envision or BMG CLARIOstar plate reader following incubation at room temperature for 10 minutes to measure cell viability. Raw data from each test well was normalized to the average (Avg) signal of non-infected cells (Avg Cells; 100% inhibition) and virus infected cells only (Avg Virus; 0% inhibition) to calculate % inhibition of CPE using the following formula: % inhibition = 100*(Test Cmpd – Avg Virus)/(Avg Cells - Avg Virus). The SARS CPE assay was conducted in BSL-3 containment with plates being sealed with a clear cover and surface decontaminated prior to luminescence reading. Reference compounds for CPE assay were made available by SRI.

##### Method for measuring cytotoxic effect of CLQ, CLBQ14 and CLCQ

Compound cytotoxicity was assessed in a BSL-2 counter screen as follows using the Cell Titer-Glo Luminescent Cell Viability Assay^60^. Host cells in media were added in 25μl aliquots (4000 cells/well) to each well of assay ready plates prepared with test compounds as above. Cells only (100% viability) and cells treated with hyamine at 100μM final concentration (0% viability) serve as the high and low signal controls, respectively, for cytotoxic effect in the assay. DMSO was maintained at a constant concentration for all wells (0.3%) as dictated by the dilution factor of stock test compound concentrations. After incubating plates at 37°C/5%CO2 and 90% humidity for 72 hours, 30μl CellTiter Glo (CTG) (G7573, Promega) was added to each well. Luminescence was read using a BMG CLARIOstar plate reader following incubation at room temperature for 10 minutes to measure cell viability.

#### Biochemical Screening Assays

#### ACE2 Inhibitor Screening Assay

An ACE2 Inhibitor screening assay kit with fluorogenic substrate (Catalogue #79923) was purchased from BPS Bioscience (San Diego, CA), and adapted to measure the exopeptidase activity of ACE2 in the presence and absence of inhibitors. The Fluorescence assay was performed using a black flat-bottom 96-well plate with a final reaction volume of 50 μL following the manufacturer’s instructions. We prepared 10mM stock solutions of the compounds in Dimethyl sulfoxide (DMSO). Next, we serially diluted the compounds in DMSO as follows: 100, 50, 10, 1, 0.5, and 0.1 μM for CLQ and CLBQ14; as well as 10 μM, and 1 μM for CLCQ. All experiments were performed in triplicates. Each plate contained a positive control of enzyme-treated with vehicle alone (2% DMSO), and a blank control with no enzyme. Briefly, each reaction contained 24 μL of purified recombinant human ACE2 protein (0.42ng/μL) in ACE2 buffer, 1 μL of compound at serially diluted concentrations, and 25 μL ACE2 fluorogenic substrate. The total reaction volume was 50 μL. The reaction mixtures were protected from light and incubated for 2.5 hrs at room temperature (22°C). Thereafter, the fluorescence intensities (λ_Excitation_ = 535nm, λ_Emission_ = 595nm) were measured using a Beckman Coulter DTX880 multimode plate reader. A similar experiment was conducted to measure and compare the exopeptidase activity of ACE2, in the presence and absence of Zinc Chloride (ZnCl_2_) alone, CLBQ14 alone and ZnCl_2_ in combination with CLBQ14 at concentrations ranging from 100μM to 100nM. ZnCl_2_ was serially diluted in water, and a positive control of enzyme-treated with vehicle alone (water for ZnCl_2_ only; DMSO for CLBQ14 alone; and water plus DMSO for ZnCl_2_ and CLBQ14) was carried out for this experiment. The background hydrolysis was subtracted and the data was fitted to a four-parameter logistic (variable slope) equation using GraphPad prism software 8.4.3.

#### The ACE2-Spike (RBD) Protein Interaction assay

A Spike-ACE2 binding assay kit (Cat # CoV-SACE2-1, Lot# 062320 7066) was purchased from RayBiotech (Norcross, GA). The *in vitro* enzyme-linked immunoabsorbent assay (ELISA), was and adapted and performed in a transparent flat-bottom 96-well plate. We prepared 10mM stock solutions of the compounds in Dimethyl sulfoxide (DMSO), with serially diluted the compounds in DMSO as follows: 100, 50, 10, 5, 1, 0.5, and 0.1 μM for CLQ, CLBQ14 and CLCQ. All experiments were performed in triplicates. Each plate contained positive controls (1% DMSO) and blank controls with no ACE2. Briefly, 1 μL of serially diluted compounds were incubated with recombinant SARS-CoV-2 Spike receptor binding domain (RBD) protein, pre-coated on the 96 well plates in 49 μL of 1X assay diluent buffer for 31 mins, at room temperature (22°C) with shaking at 180rpm. Next, we added 50 μL of ACE2 protein in 1X assay diluent buffer into the 96 well plate, and incubated for 2.5 hrs at room temperature (22°C) with shaking at 180rpm. Thereafter, the solution was discarded and the plate was washed consecutively four times with 300 μL 1X wash buffer, followed by the addition of the detection antibody (anti-ACE2 goat antibody). The reaction was allowed to go on for 1 hr at room temperature (22°C) with shaking at 180rpm. Then, the solution was discarded and the wash step was repeated as described above. Next, the HRP-conjugated anti-goat IgG was added to each well, and the reaction plate was further incubated for 1 hr at room temperature (22°C) with shaking at 180rpm. Again, the solution was discarded and the wash step was repeated as described above. Then, 100μL of 3,3’,5,5’-tetramethylbenzidine (TMB) one-step substrate was added to each well, and reaction mixtures were incubated in the dark at room temperature (22°C) with shaking at 180rpm for an additional 30 mins and then stopped by the addition of 50μL stop solution. The absorbance was read at 405 nm using a Beckman Coulter DTX880 multimode plate reader. The background hydrolysis was subtracted and the data was fitted to a special bell-shaped dose-response curve equation using GraphPad prism software 8.4.3.

### RESULTS

#### Efficacy of Clioquinol (CLQ) and Analogues against SARS-CoV-2 infection induced Cytopathic Effect (CPE) in Vero E6 cells

In our efforts to identify inhibitors of SARS-CoV-2 infection for potential treatment of COVID-19, we evaluated the *in vitro* antiviral activity of CLQ, and two of its derivatives, CLBQ14 and CLCQ, using a standard luminescent-based high-throughput screening (HTS) platform^59,60^ for SARS-CoV-2 infection induced CPE in African Green Monkey Kidney Vero E6 cells. We found that all three compounds inhibited SARS-CoV-2 infection induced CPE *in vitro* with 50% Inhibitory Concentration (IC_50_) values in the low micromolar concentration (Figure 1). Amongst all three analogues tested, CLQ displayed the most potent antiviral activity in the CPE assay (Figure 1). Compared to its counterparts, CLBQ14 exhibited the highest maximum inhibition at about 102.96% inhibition at 30μM (Table 1). In addition, we compared the antiviral effects of CLBQ14 and its analogues with five other known inhibitors of SARS-CoV-2 *in vitro*: Chloroquine, Hydroxychloroquine, Remdesivir, Aloxistatin and Calpain Inhibitor IV. The dose-response curves of the CLQ, CLBQ14, CLCQ and the reference compounds mentioned above were determined at multiplicities of infection (MOI) of about 0.002. We found that the IC_50_ for CLQ (12.62 μM), and its analogues [(CLBQ14, 14.69 μM) and (CLCQ,16.30 μM)] were slightly lower than the IC_50_ of Aloxistatin (16.72μM); but moderately higher than Chloroquine (1.10μM), Hydroxychloroquine (5.04μM), Remdesivir (4.42μM), and Calpain Inhibitor IV (0.41μM) (Table 2). These results suggest a potential new mechanism of action for CLQ and its congeners. Notably, this is the first report to our knowledge, revealing that CLQ and its analogues effectively inhibit the novel SARS-CoV-2 infection induced CPE.

**Figure 1.**
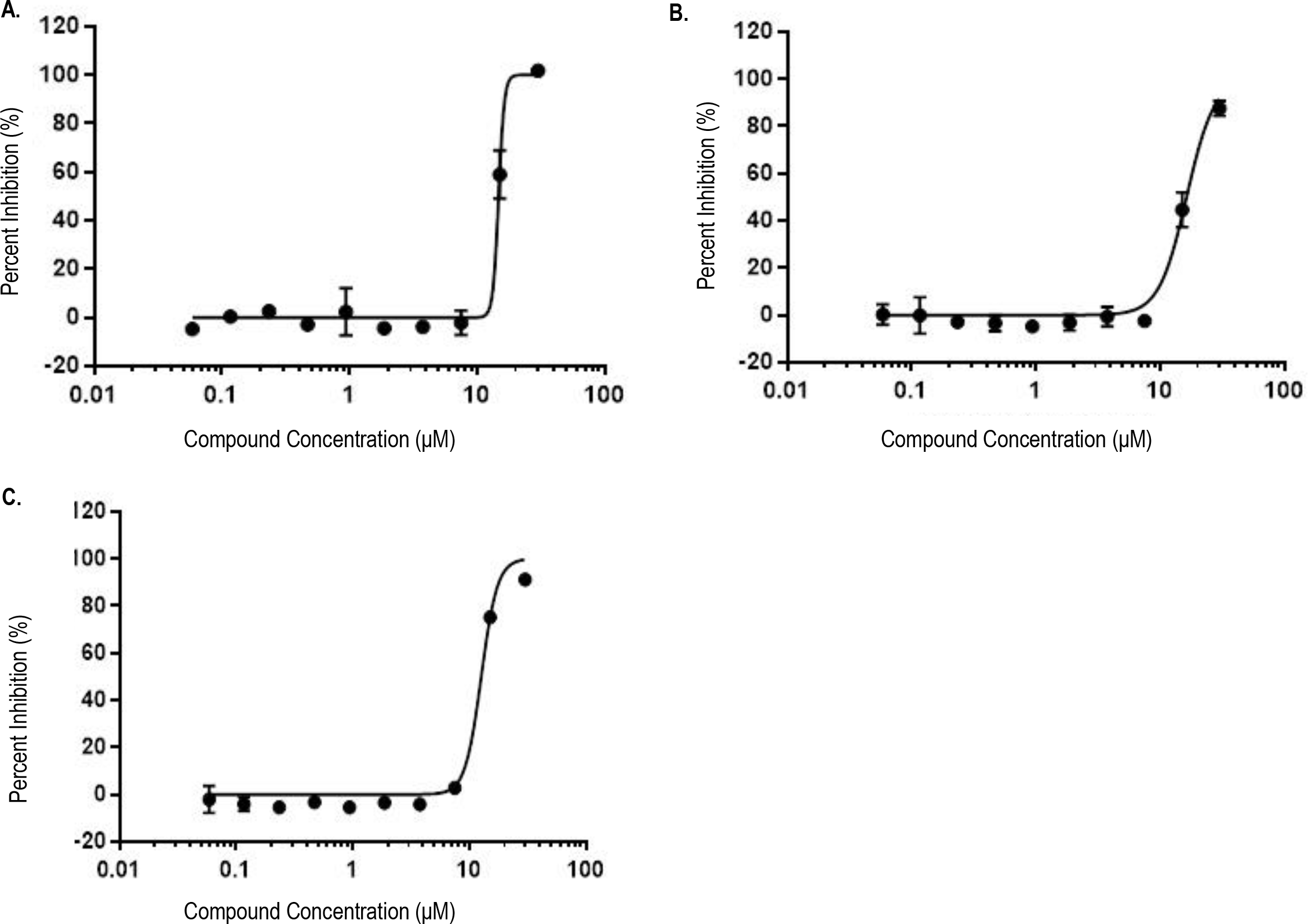
Efficacy of Clioquinol (CLQ) and Analogues against SARS-CoV-2 induced Cytopathic Effect (CPE) in Vero E6 cells: A. CLBQ14, B. CLCQ, and C. CLQ.

**Figure 2.**
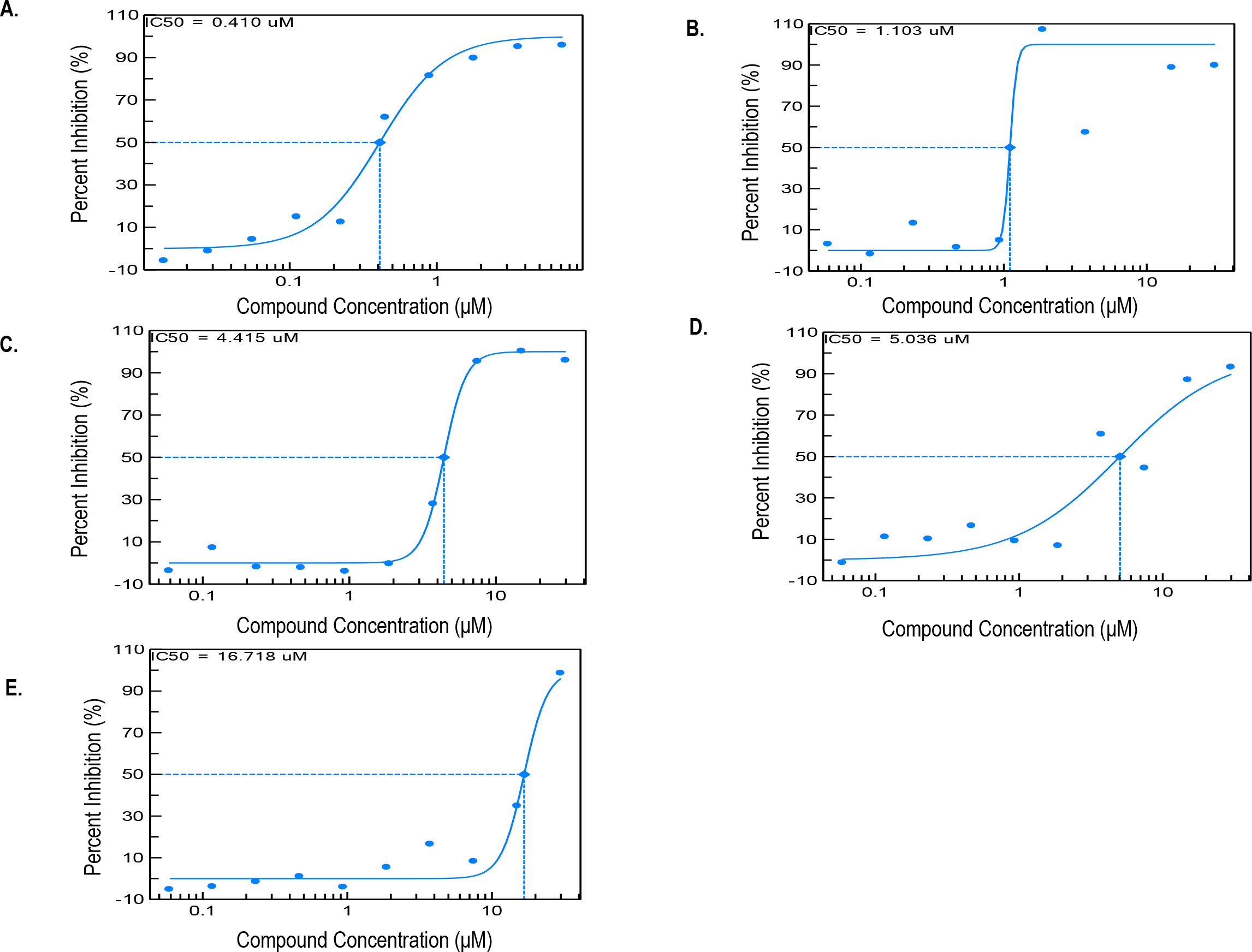
Efficacy of Reference Inhibitors against SARS-CoV-2 induced Cytopathic Effect (CPE) in Vero E6 cells: A. CalpainInhibitorIV, B. Chloroquine, C Remdesivir, D. Hydroxychloroquine, and E. E64d (Aloxistatin).

**Table 1.**
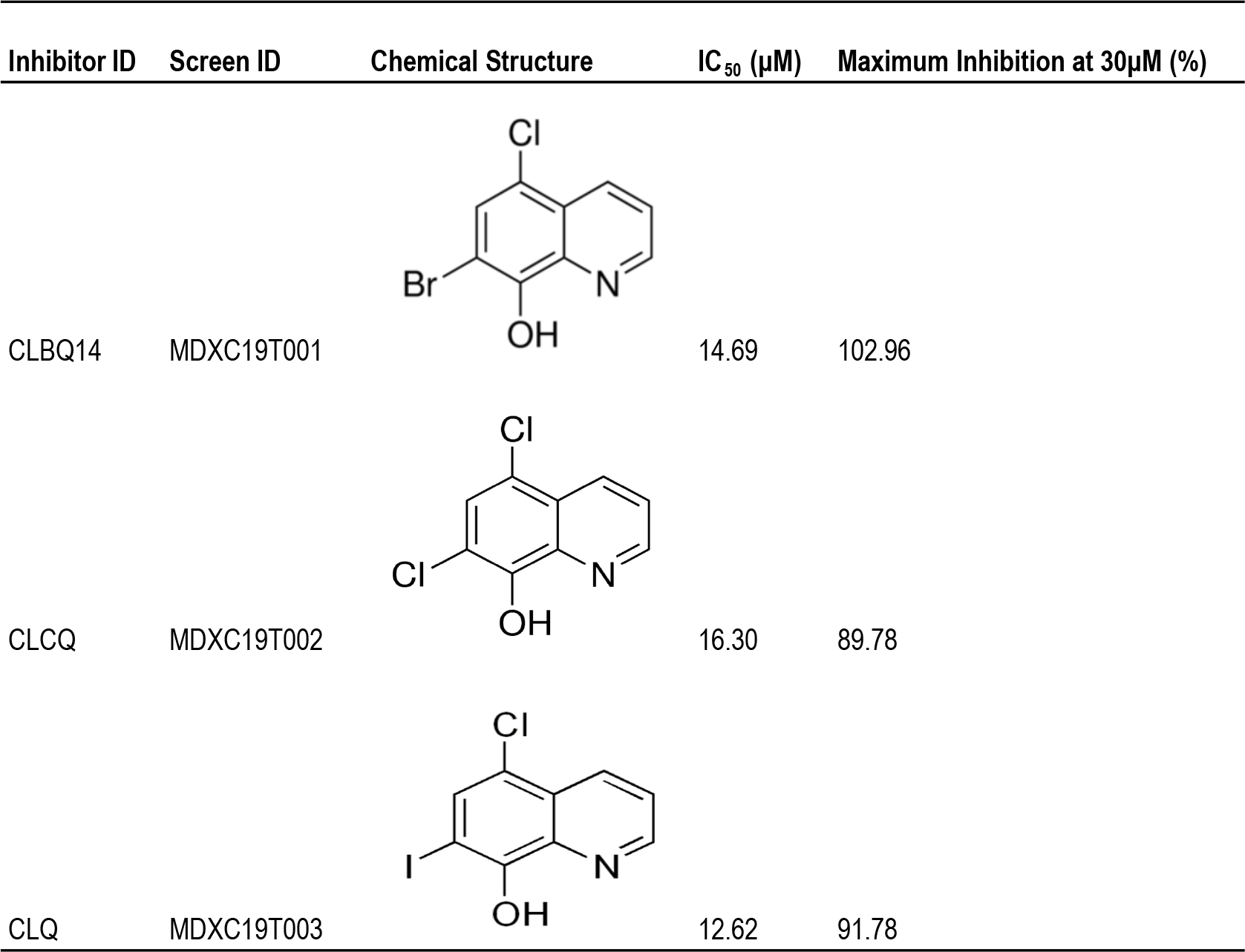
Chemical Structure and Activity of Clioquinol (CLQ) and Analogues against SARS-CoV-2 induced Cytopathic Effect (CPE) in Vero E6 Cells.

| Inhibitor ID | Screen ID | Chemical Structure | IC<sub>50</sub> (μM) | Maximum Inhibition at 30μM (%) |
| --- | --- | --- | --- | --- |
| CLBQ14 | MDXC19T001 |  | 14.69 | 102.96 |
| CLCQ | MDXC19T002 |  | 16.30 | 89.78 |
| CLQ | MDXC19T003 |  | 12.62 | 91.78 |

**Table 2.**
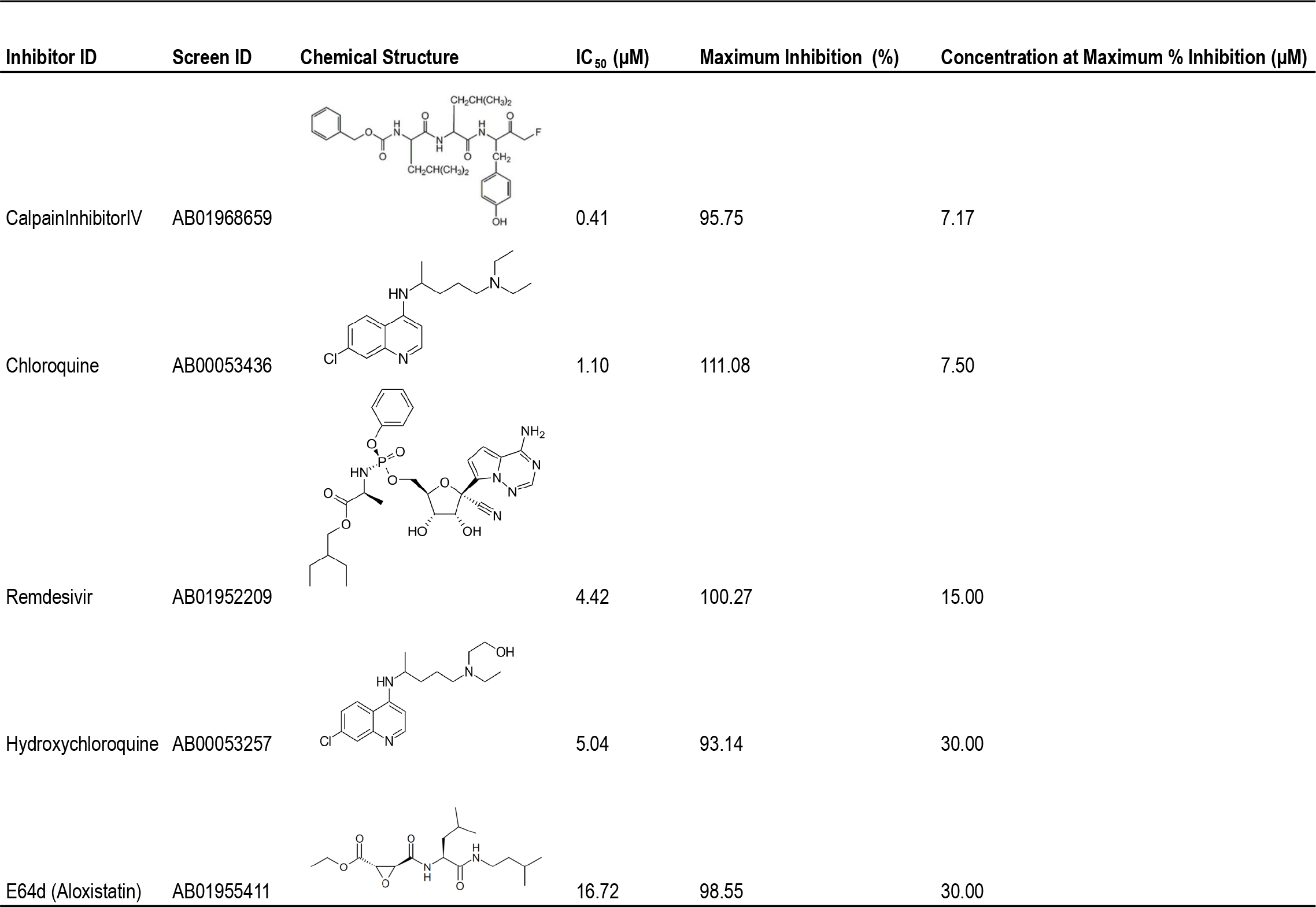
Chemical Structure and Activity of Reference Inhibitors against SARS-CoV-2 induced Cytopathic Effect (CPE) in Vero E6 Cells.

| Inhibitor ID | Screen ID | Chemical Structure | IC<sub>50</sub> (μM) | Maximum Inhibition (%) | Concentration at Maximum % Inhibition (μM) |
| --- | --- | --- | --- | --- | --- |
| CalpainInhibitorIV | AB01968659 |  | 0.41 | 95.75 | 7.17 |
| Chloroquine | AB00053436 |  | 1.10 | 111.08 | 7.50 |
| Remdesivir | AB01952209 |  | 4.42 | 100.27 | 15.00 |
| Hydroxychloroquine | AB00053257 |  | 5.04 | 93.14 | 30.00 |
| E64d (Aloxistatin) | AB01955411 |  | 16.72 | 98.55 | 30.00 |

#### Cytotoxicity Effects of CLQ and Analogues in Vero E6 cells

We determined the preliminary cytotoxicity of CLQ and its analogues (CLBQ14 and CLCQ), using a Cell Titer-Glo Luminescent Cell Viability Assay^60^. We assessed the cytotoxic effects of the various compounds in Vero E6 cells and observed that, the 50% cytotoxic concentration (CC_50_) of CLQ and its derivatives were all greater than 30 μM. However, in comparison to the other reference compounds tested, CLQ and its analogues displayed lower percent minimum viability at higher concentrations. On the other hand, we observed similar percent maximum viability for CLQ pharmacophore and the other reference compounds at lower concentrations (Table 3). This suggests that, the cytotoxic effects may not be a concern at lower concentrations of CLQ and its analogues. Additional concentrations need to be tested in future studies to determine the actual CC50 value (Table 3).

**Table 3.** Cytotoxicity of Clioquinol (CLQ) and Analogues in Vero E6 Cells, in Comparison to Reference Inhibitors of SARS-CoV-2.

| Inhibitor ID | Cytotoxicity CC<sub>50</sub> (μM) | Minimum Viability (%) | Concentration at Minimum % Viability (μM) | Maximum Viability (%) | Concentration at Maximum % Viability (μM) |
| --- | --- | --- | --- | --- | --- |
| CLBQ14 | >30.00 | 53.50 | 15.00 | 107.88 | 0.12 |
| CLCQ | >30.00 | 60.82 | 15.00 | 101.51 | 0.06 |
| CLQ | >30.00 | 61.83 | 30.00 | 105.82 | 0.23 |
| CalpainInhibitorIV | >7.17 | 98.29 | 3.59 | 104.99 | 0.22 |
| Chloroquine | >30.00 | 95.63 | 15.00 | 106.60 | 0.06 |
| Remdesivir | >30.00 | 97.49 | 3.75 | 104.49 | 0.06 |
| Hydroxychloroquine | >30.00 | 96.88 | 3.75 | 103.65 | 0.06 |
| E64d (Aloxistatin) | >30.00 | 100.06 | 15.00 | 112.76 | 0.12 |

#### Effects of CLQ and its Analogues on rhACE2 Exopeptidase Activity

We determined the effect of CLQ, CLBQ14 and CLCQ on the exopeptidase activity of rhACE2 using an adapted fluorometric assay (https://bpsbioscience.com/pub/media/wysiwyg/79923.pdf). We found that all three compounds inhibited rhACE2 activity with similar IC_50_ values in the low micromolar concentration, with CLQ being the most potent amongst all three analogues tested, at IC_50_ of 5.36μM (Table 4). To our knowledge, these results revealed for the first time that, rhACE2 is a biochemical target of CLQ and its analogues. Because, the known metal cofactor for ACE2 is Zinc^48, 61^, using the same fluorometric assay described above in the methods section, we further assessed the exopeptidase activity of rhACE2, in the presence of Zinc Chloride (ZnCl_2_) alone, CLBQ14 alone and ZnCl_2_ in combination with CLBQ14 at concentrations ranging from 100μM to 100nM. In the presence of ZnCl_2_ alone, rhACE2 displayed increasing exopeptidase activity. On the other hand, in the presence of ZnCl_2_ in combination with CLBQ14, we observed an increased shift in IC_50_ value by over 28 fold compared to CLBQ14 alone (Figure 3). Interestingly, this data reveals that increasing concentrations of ZnCl_2_, titrates the inhibitory effect of CLBQ14 on rhACE2 from concentrations ranging from above 5 - 10μM, consistent with previous reports of the required optimal concentration range of Zinc for the exopeptidase activity of ACE2^61^. Taken together, these preliminary results reveal a new pharmacologic mode of action and novel target for CLQ and its analogues.

**Figure 3.**
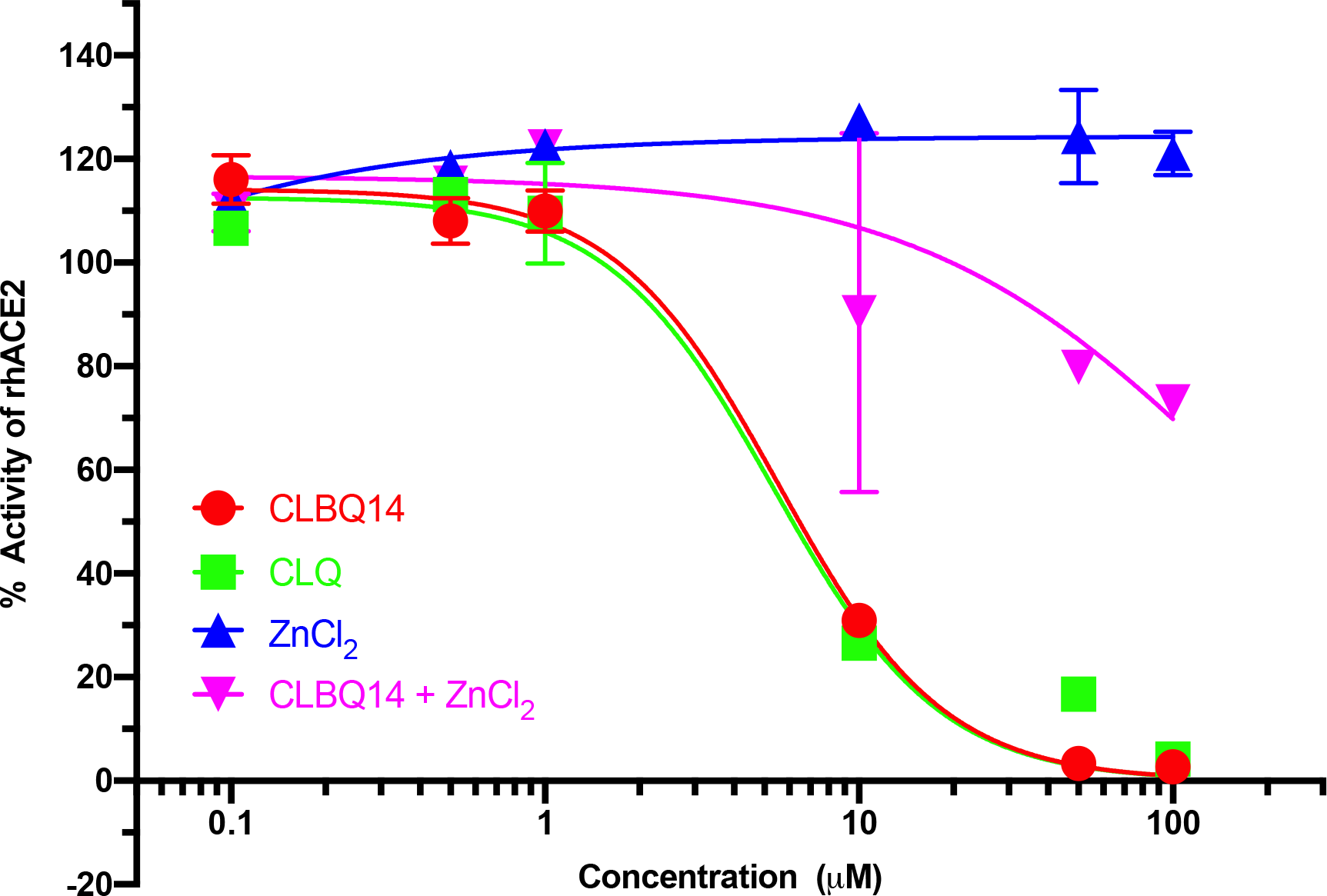
Effect of Clioquinol (CLQ) and Analogues against ACE2 Exopeptidase Activity: A. CLBQ14 (Circles - red), B. CLQ (Squares - green), and C. ZnCl_2_ (Triangle – blue), and D. CLBQ14 and ZnCl_2_ (Inverted Triangles – magenta).

**Table 4.** Activity of Clioquinol (CLQ) and Analogues against ACE2 Exopeptidase Activity and ACE2 and SARS-CoV-2 Spike (RBD) Protein Interaction.

| Inhibitor ID | IC<sub>50</sub> (μM) |  |  |
| --- | --- | --- | --- |
|  | ACE2 Exopeptidase Activity Assay | Spike (RBD)-ACE2 Interaction Assay (IC<sub>50.1</sub> (μM)) | Spike (RBD)-ACE2 Interaction Assay (IC<sub>50.2</sub> (μM)) |
| CLBQ14 | 5.55 | 2.76 | 3.06 |
| CLCQ | <10 | 1.74 | 1.91 |
| CLQ | 5.36 | 0.85 | 18.15 |
| *CLBQ14 and ZnCl<sub>2</sub> | 159.00 | ND | ND |
| * ZnCl<sub>2</sub> | ND | ND | ND |

\*Higher Concentrations need to be conducted to determine IC<sub>50</sub>

#### Effects of CLQ and its Analogues on rhACE2 and Spike (RBD) Protein Interaction

The interaction of human ACE2 receptor with SARS-CoV-2’s Spike protein receptor binding domain is a critical first step in the process required for viral entry into host cells^40–43^. Using an adapted *in vitro* enzyme-linked immunoabsorbent assay (ELISA) (https://doc.raybiotech.com/pdf/Manual/CoV-SACE2_2020.07.09.pdf), we evaluated the effect of CLQ, CLBQ14 and CLCQ on the binding affinity of rhACE2 and RBD of S protein at concentrations ranging from 100 μM to 100 nM. Surprisingly, we observed a unique bell shaped dose-response curve for all three compounds with higher inhibition of ACE2-Spike (RBD) protein interaction at lower compound concentrations compared to higher concentrations (Figure 4). The bell shaped curve generated two IC_50_ values (IC_50_1_ and IC_50_2_) as shown in Table 4. We found that all three compounds had similar IC_50_ values in the low micromolar concentration ranging from 0.85 μM to 2.76 μM for IC_50__1; however CLQ displayed a higher IC_50_2_ at 18.15 μM (Table 4). The unconventional dose response curve observed in this interaction assay, could be an indicator of additional binding site(s) and/or target(s), for the CLQ pharmacophore, such as other sites on ACE2 or the Spike (RBD) protein. Again, these findings are the first report to reveal that CLQ and its analogues inhibit and interfere with the binding between human ACE2 receptor and SARS-CoV-2 Spike RBD protein *in vitro*. These results suggest that the CLQ and its derivatives might be promising leads for clinical development of novel SARS-CoV-2 entry inhibitors and potential COVID-19 therapeutics.

**Figure 4.**
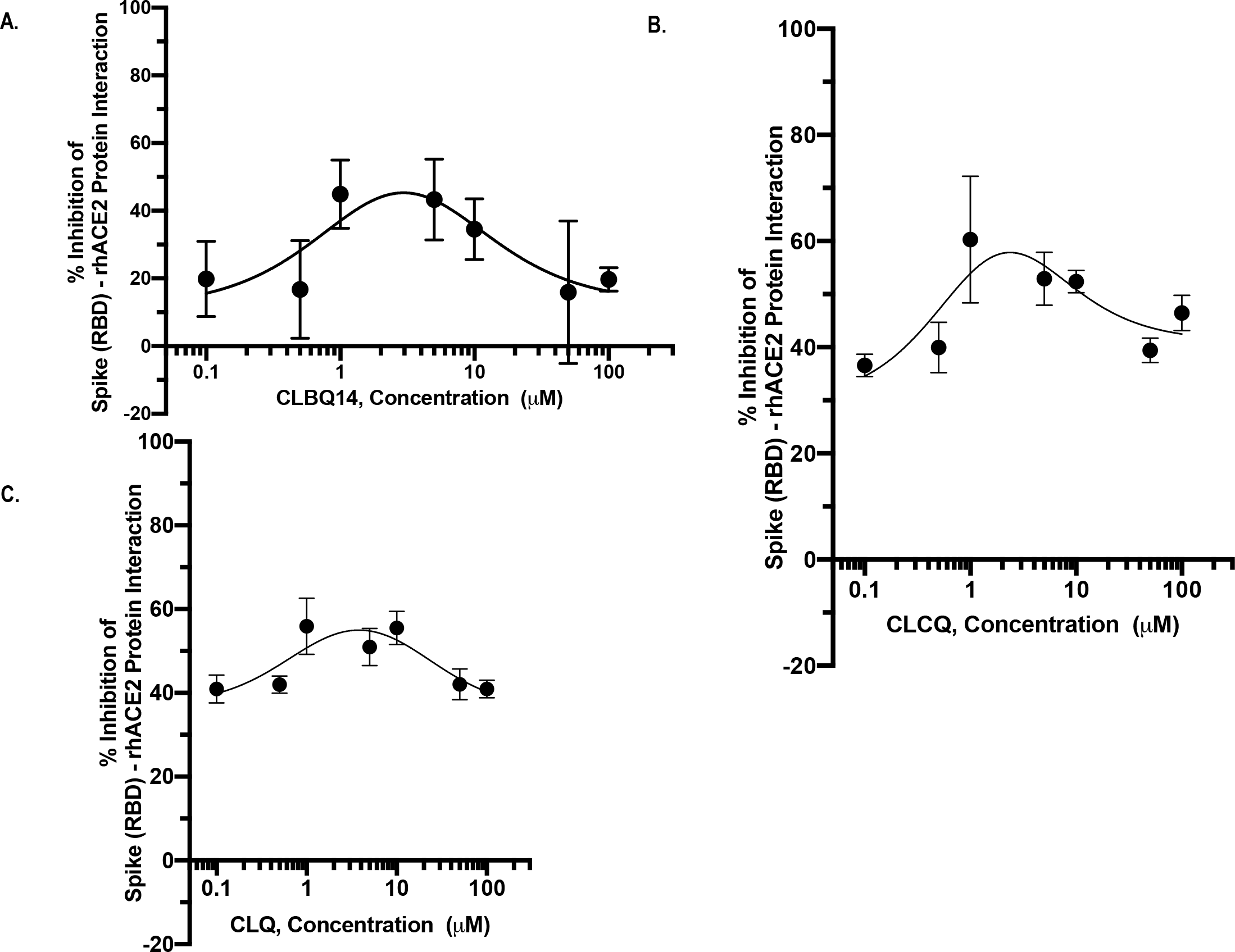
Inhibition of ACE2 and SARS-CoV-2 Spike (RBD) Protein Interaction by Clioquinol (CLQ) and Analogues: A. CLBQ14, B. CLCQ, and C. CLQ.

## DISCUSSION

Given the ongoing COVID-19 pandemic and the emerging virulence of novel SARS-CoV-2 strains, there is an urgent need to accelerate the development of effective therapeutic agents as countermeasures against this pathogen. In this study, we applied three independent approaches, to investigate the possibility of CLQ and its analogues as potential inhibitors of the SARS-CoV-2 infection *in vitro*, and gathered strong evidence that this pharmacophore are promising leads for the discovery and pre-clinical development of novel SARS-CoV-2 entry inhibitors and potential COVID-19 therapeutics. To our knowledge, this is the first report revealing rhACE2 as a novel target for CLQ and its analogues, a new pharmacologic mode of action for an old antimicrobial. Taken together, our *in vitro* findings that CLQ significantly inhibited binding of rhACE2 receptor with SARS-CoV-2 Spike (RBD) protein and SARS-CoV-2 infection induced CPE, strongly supports the notion that CLQ and its congeners could be potential drugs and/or chemical probes in the development of counter measures against viral entry into host cells.

The availability of simple, rapid, cellular high throughput screening and well-characterized biochemical assays enabled us to quickly discover novel inhibitors of SARS-CoV-2 infection *in vitro*. We successfully identified and characterized CLQ, a known metal chelator and zinc ionophore, as a novel inhibitor of SARS-CoV-2 infection induced CPE. Using two structural analogues of CLQ (CLBQ14 and CLCQ) in hand, we were able to further explore the impact of the active CLQ pharmacophore on the novel coronavirus infection, the exopeptidase activity of rhACE2 and the interaction of rhACE2 with SARS-CoV-2 Spike (RBD) protein, all critical steps/processes in the pathogenesis of COVID19. All three analogues displayed similar potent inhibition in the low micromolar range, against SARS-CoV-2 infection induced CPE, rhACE2 activity and its interaction with Spike Protein. In this study, we also compared the dose-response curves of antiviral effects of CLQ and its analogues with five other known inhibitors of SARS-CoV-2 *in vitro*: Chloroquine, Hydroxychloroquine, Remdesivir, Aloxistatin and Calpain Inhibitor IV and found that CLQ’s potency was better and comparable to Aloxistatin; but had lower efficacy than the other reference inhibitors (Table 2). It is important to note that the Vero E6 cells used for the SARS-CoV-2 infection induced CPE assay were first sorted by flow cytometry by SRI for selection of cells that had higher levels of ACE2 expression to increase the efficiency of infection. Therefore, the observed IC_50_ values may be higher than the actual IC_50_ values in cells that do not have high levels of ACE2 expression. Moreover, we observed that the IC_50_ values of the compounds in the biochemical assays were much lower than the IC_50_ in the cellular antiviral assay. We also assessed the cytotoxic effects of the compounds in Vero E6 cells and observed that CLQ and its analogues displayed lower percent minimum viability at higher concentrations compared to the other reference compounds tested. However, we observed similar percent maximum viability for CLQ pharmacophore and the other reference compounds at lower concentrations (Table 3). This suggests that, the cytotoxic effects may not be a concern at lower concentrations of CLQ and its analogues. In addition, the observed IC_50_ values for inhibition of rhACE2 exopeptidase activity and rhACE2-RBD interaction were in the low micromolar range, suggesting that we may need lower concentrations for *in vivo* activity. Furthermore, we have other preliminary cytotoxicity results from prior *in vivo* studies on CLQ and its analogues that reveal no significant toxicity at much lower concentrations below nanomolar range (unpublished data). Therefore, additional *in vivo* cytotoxicity studies for Vero E6 cells should be conducted at a wider range of concentrations.

Throughout our study, we consistently observed a correlation between the high potency of CLQ compared to its other two analogues in the antiviral screen, inhibition of rhACE2 metalloprotease activity, and its ability to disrupt the binding of rhACE2 with SARS-CoV-2 Spike (RBD) protein. Amongst all three compounds, CLQ displayed the highest potency in all three independent assays; except for IC_50_2_. Hence, validating its potential as a therapeutic option for the treatment of COVID19. Clioquinol and its derivatives belonging to 8-hydroxyquinoline structural class, have been investigated extensively in basic, translational and clinical studies because of their multiple activities as metal chelators and zinc ionophores, modulating underlying molecular and physiologic switches required for metal homeostatis *in vivo* ^14–32, 36–39^. Previously, CLQ was used to treat bacterial infections^62^; however, it was withdrawn from the clinic because of untoward effects of subacute myelo-optic neuropathy (SMON) mostly experienced in Japan in the 1950’s^29,62,63^. More recent studies revealed that SMON might be due to other biologic factors and/or pharmacogenetics primarily linked to the Japanese population^29, 62, 64^. Currently in the clinic, CLQ is approved for use in combination with other agents for treatment of inflammatory skin disorders and fungal infections in some countries^14,65^. More recently, CLQ and its newer structural derivatives have gained renewed interest as potential drugs for the development of therapeutics for neurodegenerative diseases, cancer, and infectious diseases ^14–32, 36–39^. Furthermore in previous studies, Olaleye O. et. al., serendipitously discovered CLBQ14, the bromine analogue of CLQ and characterized CLQ and additional derivatives as potent inhibitors of replicating and non-replicating *Mycobacterium tuberculosis*, using a HTS assay designed to identify novel metalloprotease inhibitors^17^. Altogether, the plethora of evidence on the broad pharmacologic spectrum of activity and metal-chelation propensity of CLQ pharmacophore, combined with its extensive clinical investigational profile, makes this structural class an attractive and promising drugs for targeting ACE2, the important zinc metalloenzyme and essential cellular receptor for SARS-CoV-2 entry into host cells^40–43^.

ACE2, a carboxypeptidase, is a known type I integral membrane protein made up of about 805 amino acids belonging to the large family of Zinc metalloproteases with high level of structural homology for a catalytic motif, containing one characteristic HEXXH + E zinc-binding consensus sequence and binding sites for inhibitor or specific substrates respectively^48^. According to earlier reports by Towler et. al., the first crystalline structures of the metallopeptidase domain of ACE2, revealed “a large inhibitor-dependent hinge bending movement of one catalytic subdomain relative to the other that brings important amino acid residues into position for catalysis,” similar to observed subdomains on other zinc metalloproteases respectively^48^. The residues critical for coordinating the binding of Zinc to ACE2 are His^374^, His^378^ and Glu^402^, according to earlier x-ray strucutres^48^. Moreover, ACE2 is activated by monovalent anions and also known to contain an inhibitor-specific anion binding site^48,61^. The reported optimal metalloprotease activity of recombinant soluble human ACE2 was found to be in the presence of 10μM ZnCl_2_^61^. This is consistent with our findings of rhACE2 exopeptidase activity assay, in the presence of Zinc Chloride (ZnCl_2_). In the presence of the newly identified potent metalloprotease inhibitors (CLQ or CLBQ14 alone), we observed a significantly decreased exopeptidase activity for ACE2 in the low micromolar concentrations (Figure 4). However, we found an increased shift in IC_50_ values when we assessed exopeptidase activity in the presence of ZnCl_2_ in combination with CLBQ14 by over 28 fold compared to CLBQ14 alone (Figure 3), suggesting that CLBQ14, might be working through zinc chelation, interaction and/or coordination. Our findings not only revealed a novel target (rhACE2) and mechanism of action for the CLQ pharmacophore; but also provides insight into potential reversibility of inhibition and one or more probable mode(s) of inhibition: 1) The concentration of CLBQ14 is titrated with excess ZnCl_2_, thus pre-occupied and unavailable to inhibit rhACE2 exopeptidase activity; and/or 2) potential competition for the similar binding sites on rhACE2. Additional mechanistic kinetic studies will be required to ascertain this notion.

Moreover as mentioned earlier, ACE2, plays an essential role in the regulation of cardiovascular and respiratory physiology ^47,48^. Its characterization as the functional host receptor for entry of the novel SARS-CoV-2 into human cells^40–43^, has raised concerns about the potential impact of newly discovered ACE2 inhibitors on cardiovascular and respiratory physiology^66–68^. Recent studies have also shown that ACE2 plays a key role in protecting the lungs from ARDS^67,68^, a severe complication of COVID-19 disease^4^. Therefore, one has to proceed cautiously when targeting ACE2^66^; without permanently inactivating its exopeptidase or other cellular functions, to avoid potential adverse effects to heart and/or lung function. Our lead compound CLQ is a weak metal chelator and zinc ionophore, that can shuttle free zinc across the membrane^31,69^. Because of these properties, CLQ may temporarily or reversibly affect ACE2 function and prevent its interaction with SARS-CoV-2 RBD protein; without permanently inhibiting its essential exopeptidase function. Because rhACE2 is a novel host target for CLQ and its analogues, the potential effect of CLQ inhibition on heart and lung function needs to be further explored *in vivo* and pre-clinical studies.

The crystal structure of full length human ACE2 revealed that the RBD on SARS-CoV-2 S1 binds directly to the metallopeptidase domain (MPD) of ACE2 receptor^40,41^, that consists of amino acid residues that coordinates zinc, providing further support for the utility of zinc chelators and/or ionophores such as CLQ and its congeners, as promising inhibitors of interaction and viral entry inhibitors. Using a sensitive ELISA, we found that CLQ and its analogues potently disrupt the interaction of ACE2 and Spike (RBD) protein, with CLQ being the most potent. Thus, supporting our findings showing that CLQ and its derivatives binds to ACE2 and inhibits exopeptidase activity. Interestingly unlike the CLQ pharmacopore, other studies revealed that (S,S)-2-{1-Carboxy-2-[3-(3,5-dichloro-benzyl)-3H-imidazol-4-yl]-ethylamino}-4-methyl-pentanoic acid (MLN-4760), a known potent inhibitor of ACE2 exopeptidase activity, belonging to a different chemical class, does not disrupt ACE2-Spike interaction in coronaviruses, SARS-CoV, SARS-CoV2, and NL63S ^70,71^ as its binding site on ACE2 is different than the site where RBD interacts with ACE2 ^48,70,72^. However, CLQ seems to affect ACE2 by reversibly chelating its zinc ion which is essential for ACE2 activity, as well as interfere with ACE2-RBD interaction. Although zinc is essential for stabilizing protein structures and altering the substrate affinity of different metalloproteins^32,73^, the effects of zinc chelation on molecular structure of ACE2 and its effects on its binding to the SARS-CoV-2 remains to be tested. Moreover, earlier molecular and structural studies also revealed that mutations in the catalytic site required for exopeptidase activity of ACE2, had no effect on Spike RBD binding to ACE2^48^. Howbeit, the unconventional dose-response bell shaped curve that we observed in our studies suggests that their may be additional binding sites and/or modes of action for CLQ and its congeners, resulting in the potent inhibition of interaction at lower micromolar concentrations; compared to higher concentrations. Although, CLQ was found to be the most potent amongst all 3 analogs, except for IC_50_2_, preliminary SAR revealed that the other two derivatives are comparable to CLQ, as they both show potent inhibition of rhACE2-RBD interaction, as well as inhibition of antiviral and anti-rhACE2 activity. Therefore, providing alternative analogues that might not have the same adverse effects experienced with CLQ in the past, potentially alleviating some of the concerns with CLQ. Additional biochemical and structural studies are required to explore other possible mechanisms of action such as competition or interaction of CLQ with RBD for binding to MPD of ACE2, thereby preventing zinc chelation and ionophore activity of CLQ. Future X-ray structures could help to better understand mode of inhibition of this pharmacophore and rational design of more potent drugs.

The strengths of our study includes, the use of a rapid multi-prong approach via three sensitive independent assays, to identify and characterize an existing clinical drug as a novel inhibitor of SARS-CoV-2 infection *in vitro*. In addition, the availability of structural analogues of CLQ, made possible a preliminary structure activity relationship studies (SAR), which revealed similarity between the IC_50_ values of CLQ and its structural analogues. However, our study has some limitations such as the use of Vero E6 cells that were selected for high expression of ACE2 in the antiviral assay, a HTS designed to rapidly screen for inhibitors of infection induced CPE. An additional limitation is that, the amount of zinc in the purified rhACE2 supplied from BPS and RayBiotech assays were unknown. Future metal dependent studies with apoenzymes will be required to determine the amount of zinc. Considering that CLQ is a known zinc chelator and ionophore, an understanding of the physiologic amount of zinc required for inhibition will be critical for optimal efficacy. Therefore, for these two limitations, the measured IC_50_ values for the compounds may not be representative of the actual *in vitro* IC_50_, which may be lower. However, the remarkable consistency in the observed strong correlation between CLQ and its congener’s antiviral activity, *in vitro* rhACE2 inhibition and disruption of ACE2-RBD protein interaction, reduces these concerns.

## CONCLUSION AND SIGNIFICANCE

The impact of the COVID-19 pandemic on human health, healthcare systems, and the global economy^74^ has imposed an urgent call/pressing need for the development of novel antivirals. Rapid clinical development of anti-COVID19 treatments could be accelerated by discovery of re-purposed clinically approved drugs with new mechanisms of action and/or multiple cellular targets that could potentially disrupt viral pathogenesis/survival and/or prevent the viral entry/interaction with host receptor, ACE2. The body of evidence on the broad pharmacologic spectrum of activity, metal-chelation propensity and zinc ionophore activity of CLQ pharmacophore, combined with its extensive clinical investigational profile, makes this structural class, attractive and promising drugs for targeting rhACE2. Using a multi-prong approach, we discovered and characterized CLQ, a clinical drug and two of its analogues (CLBQ14 and CLCQ) as potent inhibitors of SARS-CoV-2 infection induced CPE *in vitro*; rhACE2 metalloprotease activity; and the binding of rhACE2 with SARS-CoV-2 Spike (RBD) protein. Altogether, these novel findings provide insights into a new mode of action and molecular target(s) for CLQ and its derivatives. Thus, validating this structural class as promising leads for clinical development of novel SARS-CoV-2 entry inhibitors and potential COVID-19 therapeutics. Because rhACE2 is a host target, it reduces the concerns for development of drug resistance, which is usually seen with drugs that target viral genes. Further SAR, computational/molecular modeling and X-crystal structure studies will aid the rational design and synthesis of more potent inhibitors in the CLQ-containing, 8-hydroxylquinoline structural class. Our studies not only provides an additional new drug class with zinc chelating and ionophore properties, in the pipeline for urgent quest for therapeutic management for anti-COVID19, but also suggests that there could be the potential physiologic relevance of zinc homeostatis in SARS-CoV-2 infection and COVID19 pathogenesis. In addition, CLQ and its derivatives could be used as chemical probes to study the biology of host-pathogen interaction in the context of SARS-CoV-2 infections. In the future, the functional importance of molecular and cellular regulation of host and viral zinc dependent genes/proteins in SARS-CoV-2 pathogenesis and survival may be better understood and targeted with available zinc chelators, ionophores, and transporters. Moreover, unlike MLN-4670 another known ACE2 inhibitor^48,70,71^, our results not only show that CLQ and its analogues inhibits rhACE2, with antiviral activity, but also suggests that CLQ pharmacophore, potently disrupts the interaction of rhACE2 and Spike (RBD) protein. To this end, we provide strong cellular and biochemical evidence supporting the notion that CLQ, CLBQ14 and CLCQ, could serve as a potential lead series for the pre-clinical development of new anti-COVID19 treatments. The expectation is that the development of new anti-COVID19 treatments with dual activity against viral and host entry target could help combat the issue of emerging drug resistant strains, drug-drug interactions, reduction in the cost of treatment, possibly increase patient compliance and improve patient care as well as reduce the mortality rate due to SARS-CoV-2 infection. Therefore, we propose pharmacologic and clinical studies to further explore CLQ and/or its derivatives as treatment options in the tool box for combating this novel coronavirus, and be evaluated in conjunction with other available therapeutics to reduce COVID19 morbidity and mortality as well as potential drug to drug interactions^75–78^ encountered with other drugs.

## Author Contributions

OAO conceived the study and performed biochemical experiments (rhACE2 inhibitor screening assay and rhACE2-Spike (RBD) protein interaction experiments); OAO and MK performed experimental design, data analysis and interpretations. OAO, MK, CO and TA wrote the manuscript.

